# Lymph flow directs rapid neutrophil positioning in the lymph node in infection

**DOI:** 10.1101/2020.09.20.302075

**Authors:** Jingna Xue, Yujia Lin, Darellynn Oo, Jianbo Zhang, Flavia Jesus, Ava Zardynezhad, Luiz G. N. de Almeida, Daniel Young, Antoine Dufour, Shan Liao

## Abstract

Soon after *Staphylococcus aureus* (*S. aureus*) skin infection, neutrophils infiltrate the LN via the high endothelial venules (HEVs) to restrain and kill the invading microbes to prevent systemic spread of microbes. In this study, we found that rapid neutrophil migration depends on lymph flow, through which inflammatory chemokines/cytokines produced in the infected tissue are transported to the LN. Without lymph flow, bacteria accumulation in the LN was insufficient to stimulate chemokine production or neutrophil migration. Oxazolone (OX)-induced skin inflammation impaired lymphatic function, and reduced chemokines in the LN after a secondary infection with *S. aureus*. Due to LN reconstruction and impaired conduit-mediated lymph flow, neutrophil preferentially transmigrated in HEVs located in the medullary sinus, where the HEVs remained exposed to lymph-borne chemokines. Altered neutrophil migration resulted in persistent infection in the LN. Our studies showed that lymph flow directed chemokine dispersal in the LN and ensured rapid neutrophil migration for timely immune protection in infection. The impaired lymph flow and neutrophil migration may contribute to the frequent infection in skin inflammation, such as atopic dermatitis.

## Introduction

Skin and soft tissue infections (SSTIs) are the most frequent microbial infections in humans. Infection occurs when the skin barrier is broken by wounds or skin diseases. Following a skin infection, some microbes may travel through the lymphatic vessels to the draining lymph node (LN) (1–4). Neutrophils start entering the LN via the high endothelial venules (HEVs) within an hour to restrain and kill the invading microbes and to prevent systemic spread of microbes (5–8). HEVs are located deeply in the T cell zones of LNs (9–11). Yet, how HEVs can react rapidly to skin infection is not unclear. Skin inflammation, such as atopic dermatitis (AD), are often associated with recurrent Methicillin-resistant *Staphylococcus aureus* (MRSA) infection (12). These antibiotic resistant infections are tough to treat because of lacking understanding of the mechanism.

Lymph flow brings microbes and soluble factors (free-form antigens, chemokines and cytokines) from the infected tissue into the LN soon after an infection. Large molecules and microbes in the afferent lymph are usually captured by CD169^+^ subcapsular sinus (SCS) macrophages (13–17), and SCS dendritic cells (DCs) (18). As such, most microbes are restricted in the SCS after a skin infection (1, 19). The activated SCS macrophages further secrete pro-inflammatory cytokines to activate/recruit other types of cells to the SCS (1, 20). Antigen diffusion exponentially decays with distance, thus cannot effectively reach the T cell zone (21). Conduit-mediated lymph flow delivers small molecular weight (MW) antigens to LN resident DCs in the T cells zone or to B cells in the B cell follicles for effective antigen recognition (22–24). LN conduits are composed of a core of extracellular matrix (ECM) proteins wrapped by fibroblastic reticular cells (FRCs). Lymph flow along the conduits delivers small MW soluble factors (MW < 70 kDa), such as chemokines and cytokines, to HEVs (26). Whether and how lymph flow regulates neutrophil migration are not unclear.

The lymph-borne factors as well as the physical flow of lymphatic fluid both heavily contribute to maintaining HEV function. Constant lymph flow is required to maintain HEV specific gene expression and LN cell homeostasis (27–29). A recent study showed that lymph flow activates a mechanosensitive ion channel, Piezo1, on FRCs which plays a role in maintaining optimal HEV function in Peyer’s Patches (30). During an immune response, LN stromal cells and conduits undergo substantial remodeling to accommodate cell expansion (31, 32). Inflammation, including oxazolone (OX)-induced skin inflammation induces LN cell expansion and reduces HEV-gene expression (33–36). The reduced HEV gene expression is associated with impaired lymphatic function during OX-skin inflammation (37, 38). It remains unclear how changes in lymphatic function during skin inflammation impact neutrophil migration if exposed to a secondary infection.

In this study, by blocking lymph flow with lymphatic vessel suture or using OX-skin inflammation-induced LN remodeling, we showed that neutrophil migration depends on conduit-mediated lymph flow, which directs inflammatory chemokine dispersal for effective communication between infected tissue and the HEVs in the LN. Compromised lymph flow resulted in interrupted neutrophil positioning and persistent infection.

## Materials and Methods

### Animals

C57BL/6 mice were purchased from the Jackson Laboratory. LysM-GFP mice were a kind gift from Dr. Paul Kubes’ lab and were bred at the Health Sciences Animal Resource Center at the University of Calgary. All experiments were performed using 6-10 weeks-old female mice. All animal protocols were reviewed and approved by the University of Calgary Animal Care and Ethics Committee and conformed to the guidelines established by the Canadian Council on Animal Care.

### Suture afferent lymphatic vessel to popliteal LN

As previously described (39), mice were anesthetized with Ketamine/Xylazine. Then 5 μL 1% Evans blue in saline was injected into mouse’s footpads to visualize afferent lymphatic vessels. A small cut was made at the skin on the left leg to expose the lymphatic vessels. Afferent lymphatic vessels were sutured with 0.7 metric monofilament polypropylene. The skin incision was then closed with sutures. The contralateral right leg received the Evans blue injection and infection without suture as positive control.

### Oxazolone or FITC skin sensitization

To use inguinal LNs (iLNs), 200 µL of 4% OX in acetone (Sigma), or 2% FITC (Sigma) in 1:1 (v/v) acetone/dibutyl phthalate mixture was applied on the skin of the shaved abdomen or shaved flanks, respectively. For popliteal LNs (pLNs), 130µL 4% OX was applied on the skin of the shaved leg, or 10 µL 2% FITC was injected into the footpad.

### Methicillin-resistant *S. aureus* infection and CFU test

For iLNS, 50 µL (2.5×10^7^) *S. aureus* was intradermally injected at the right side of flank (for iLNs), or for pLNs, 20 µL (2.5×10^7^) was injected in the right footpad. LNs were collected and homogenized using a VWR Pellet Mixer (VWR) in 300 µL of sterile PBS. Tissue homogenate (100 µl) was serially diluted in PBS, then plated onto BHI agar plates and incubated at 37 °C for at least 15h after which colony-forming-units (CFU) were counted.

### Flow Cytometry

The LNs were prepared for flow cytometry by being gently pressed through a 40-µm strainer to create a single cell suspension and then washing with FACS buffer. The cells were blocked using anti-CD16/32 antibody for 5 min at room temperature, then stained for 30 min with specified conjugated markers of interest, on ice and in darkness. After washing with FACS buffer, the cells were fixed with fixation buffer (2% PFA in FACS buffer). Flow cytometry analysis was done on FlowJo.

### Immunohistochemistry Staining

Tissues were harvested from euthanized mice, embedded directly into OCT, and frozen down in dry ice. For some studies, the tissue of interest was fixed with 4% paraformaldehyde overnight and then kept in 30% sucrose for 2 hours before embedding in OCT and freezing on dry ice. 10 to 20 µm cryosections were blocked with 5% mouse serum for 1-2h. Samples were incubated overnight with primary antibodies. After three washes with PBS, samples were incubated with secondary antibodies for 60 min.

### Whole mount staining and live staining

LNs were fixed in 4% paraformaldehyde in PBS then stored in PBS. Samples were blocked with 5% mouse serum plus 0.1% Triton X-100 in PBS for 1h. Then, samples were incubated for 4h with conjugated antibodies in blocking solution. After three washes in PBS with 0.1% Triton X-100, samples were imaged by confocal microscopy.

For LN whole mount staining on SCS, LN were stained with conjugated antibodies in PBS after harvesting for 20-30 min. After three washes in PBS with 0.1% Triton X-100, samples were fixed with 4% paraformaldehyde for 20-30 min before imaging by confocal microscopy.

### Optical clearing with benzyl alcohol and benzyl benzoate (BABB)

Fixed LNs were incubated twice in 50% BABB in methanol (BABB: 1:2 mixture of benzyl alcohol and benzyl benzoate), and two to three times in 100% BABB. LNs were then kept in a microwell with 100% BABB on the slide, and coverslips were mounted on the microwell for imaging.

### Cytokines/Chemokines discovery assay

Control and OXd4 mice were infected with 2.5×10^7^ *S. aureus* in the right flank. iLNs were collected at 4hpi, and the proteins were extracted by using RIPA lysis buffer (with protease and phosphatase inhibitor). Samples were sent to Eve technology to perform the discovery assay.

### Time-lapse imaging

Mice were anesthetized with Ketamine/Xylazine. Skin and the adipose tissue around pLN were carefully removed to expose pLN. The mouse was placed on the heating pad and fixed on the stage by surgical tapes for time-lapse imaging under a multiphoton (MP) microscope. The videos were taken as soon as 10μL FITC was injected in the footpad. Time-lapse images were taken at 20 seconds/frame.

### Microscopy

Fluorescent images were taken with an SP8 multiphoton microscope (Leica). The objectives were 20× in air, 25× in water, and 63× in oil for different studies. The three-dimensional reconstruction and the diameter measurement were performed with Leica LAX software. Some of the analysis and image brightness adjustment were performed with the ImageJ software.

### Proteomic data and bioinformatics analysis

Briefly, LNs from control (uninflamed mice, n = 4) or OXd4 (n = 4) mice with *S. aureu*s at 4hpi were subjected to a shotgun proteomics analysis (Supplementary Tables 1-2). Whole sample biomasses from the biopsies were required for data collection. All liquid chromatography and mass spectrometry experiments were carried out by the Southern Alberta Mass Spectrometry (SAMS) core facility at the University of Calgary, Canada. Analysis was performed on an Orbitrap Fusion Lumos Tribrid mass spectrometer (Thermo Scientific) operated with Xcalibur (version 4.0.21.10) and coupled to a Thermo Scientific Easy-nLC (nanoflow Liquid Chromatography) 1200 system. Spectral data were matched to peptide sequences in the mouse UniProt protein database using the Andromeda algorithm (ADD ref PMID: 21254760) as implemented in the MaxQuant (ADD ref PMID:19029910) software package v.1.6.0.1, at a peptide-spectrum match FDR of < 0.01.

### Statistics

The data were expressed as mean, ± standard error (SEM). Quantitative data were tested using statistical tests in the GraphPad Prism software. Results were considered statistically significant at P < 0.05.

## Results

### Rapid neutrophil recruitment in the LN after infection depends on lymph flow

Currently, there is no genetic model to block lymph flow without impact LN microenvironment. To determine if lymph flow regulates rapid neutrophil migration in HEVs in the LN, we blocked lymph flow by suturing afferent lymphatic vessels from the footpad to the popliteal LN (pLN) as previously described (**Figure 1A**). Using FITC as a tracer, lymph could not reach HEVs when lymphatic vessels were sutured (**Figure 1B**) (40). We sutured lymphatic vessels at three conditions: a) before the infection; b) 10 min post-infection; c) 2 hour-post-infection (hpi). The contralateral LNs which were infected and injected Evans blue were collected as controls. All the LNs were collected at 4 hpi (**Figure 1C)**. Using LysM-GFP reporter mice, by whole mount imaging on the LN SCS (indicated by Evans blue dye), neutrophil accumulation was detected in the SCS in control LNs. When lymphatic vessels were sutured before the infection, few neutrophils were detected in the SCS. When lymphatic vessels were sutured at 10 min or 2 hpi, slightly more neutrophils were detected in the SCS compared to the pLNs sutured before infection, but substantially fewer than the control pLNs (**Figure 1D**). We then performed the same suture protocol using WT C57BJ/6 mice to quantify neutrophils (CD11b^+^Ly6G^+^) by flow cytometry. Consistently, neutrophil recruitment in the pLNs was significantly suppressed when lymphatic vessels were sutured before or after infection (**Figure 1E**). Cryosections of WT mice LN with anti-Ly6G and anti-Lyve-1 immunofluorescent (IF) staining further showed neutrophil recruitment was significantly suppressed when lymphatic vessels were sutured either before or short time after infection. When lymphatic vessels were sutured, most neutrophils were detected in the MS (**Figure 1F**). Collectively, these data showed that rapid neutrophil migration to the LN depends on lymph flow. When lymph was blocked, neutrophils were preferentially located in the MS at early time after infection.

**Figure 1.**
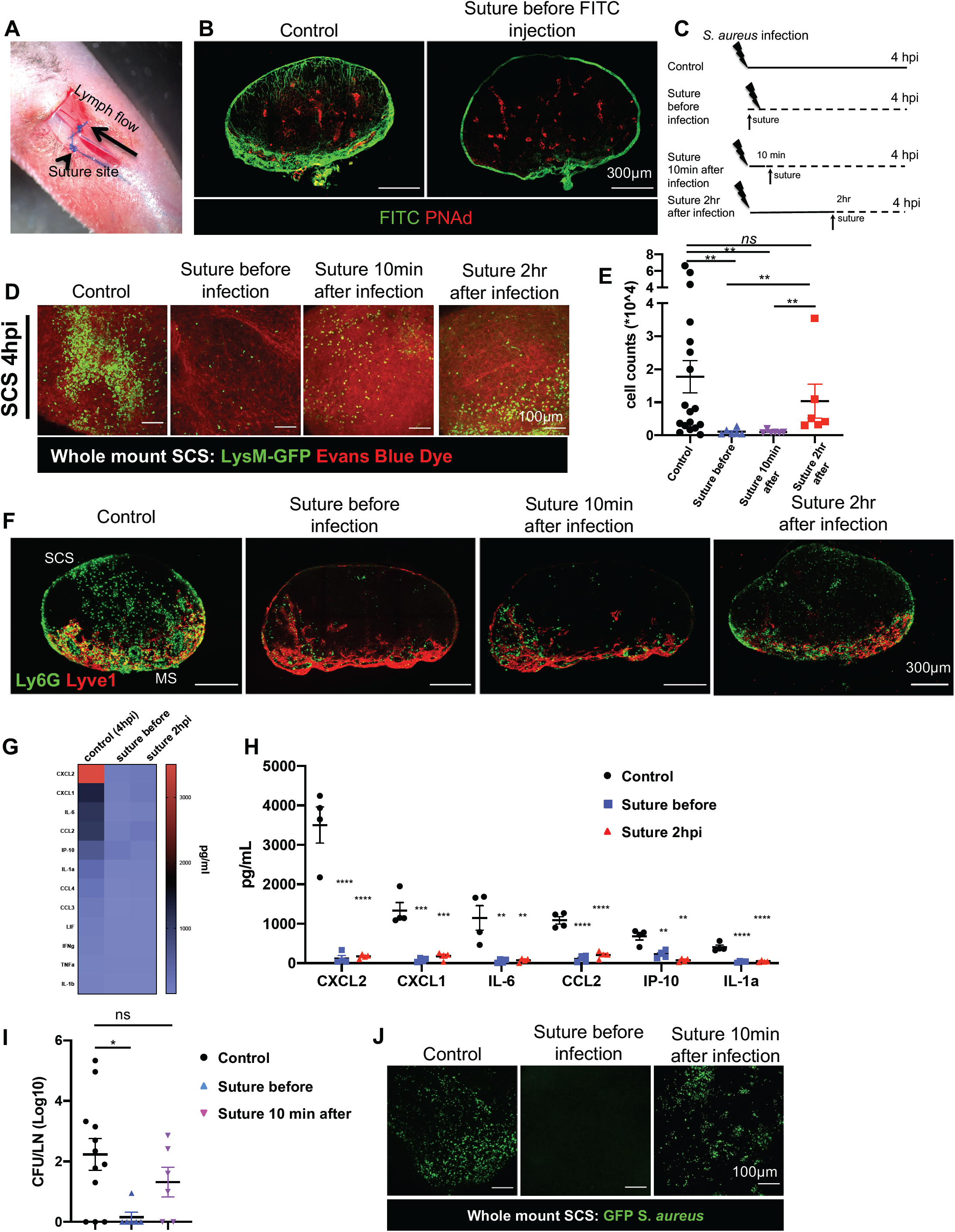
Lymph flow transports inflammatory chemokine for rapid neutrophil migration in the LN. (**A**) Representative picture of the procedure of suturing afferent lymphatic vessels. The picture shows that Evans blue dye could not pass the site where the afferent lymphatic vessels were sutured. (**B**) Control and sutured WT mice were intradermally injected with FITC at the footpad, and the pLNs were collected at 4hpi. Cryosections showed FITC could not reach PNAd^+^ HEVs in pLNs when lymphatic vessels were sutured. n=5 per group. (**C**) Schematic diagram of lymphatic suture experimental design. In the suture-before-infection group, the afferent lymphatic vessels in the left leg were sutured, then 2.5×10^7^ *S. aureus* were intradermally injected in the left footpad, and pLNs were collected at 4hpi. In suture-10 min-after-infection and suture-2h-after-infection groups, *S. aureus* were infected in the left footpad. After 10 min or 2h, the afferent lymphatic vessels were sutured and pLNs were collected at 4hpi. Evens blue dye (5 μl) were injected to visualize lymphatic vessels for the surgery. The contralateral right footpad was injected with 2.5×10^7^ *S. aureus* and Evens blue (5 μl), and the pLNs were collected at 4hpi as positive controls. (**D**) Using LysM-GFP mice to visualize neutrophils, the whole mount pLNs were imaged on the SCS (indicated by Evans blue dye (red)). Original magnification, 250×. n=3-5 per group. (**E**) Using WT C57BL/6 mice, the numbers of neutrophils (CD11b^+^Ly6G^+^) in the draining pLN were quantified by using flow cytometry. Data were mean ±SEM; Mann-Whitney test, *P<0.05; n=6. (**F**) Using WT C57BL/6 mice, cryosections of pLNs were stained with anti-Ly6G (green, neutrophils) and anti-Lyve1 (red, lymphatic endothelial cells)). n=4-5 per group. (**G**) Heatmap of cytokines/chemokines discovery array. The mice were treated, and samples were collected similar to the schematic diagram shown in Figure 1C. The draining pLNs were homogenized in lysis buffer to extract proteins. (**H**) Graph of Individual-values of several high concentration cytokines/chemokines. The concentration of other factors was provided in Table 1. Data shown as mean ±SEM, One-way ANOVA; *P<0.05; **P<0.01; ***P < 0.001; ****P < 0.0005; ns, no significance; n=4. (**I**) Bacteria load in pLNs of suture groups shown in Figure 1C. Lymphatic vessels of the pLNs were sutured before infection or 10 min after the infection. The contralateral LNs with infection and Evan blue injection served as control. All the draining LNs were collected at 4hpi for the CFU test to show the bacteria load in pLNs of different groups. Data were mean ±SEM, Brown-Forsythe and Welch ANOVA tests; *P<0.05; ns, no significance; n=5 per group. (**J**) Using WT C57BL/6 mice, control and sutured (before infection or 10 min after infection) legs were infected with GFP labeled *S. aureus*, and the pLNs were collected at 4hpi. GFP-*S*.*aureus* distribution in the SCS at different conditions was shown using whole mount pLNs. Original magnification, 250×. n=3-5 per group.

### Lymph flow transports inflammatory chemokines/cytokines from the infected tissue to the LN

Next, we set out to investigate the mechanism of how lymph flow regulates neutrophil migration. Lymphatic vessels transport cells, bacteria, and soluble factors to the LN. Skin-derived migrating DCs or neutrophils are not ready to travel via lymphatic vessels to the LN by 4hpi (6, 41). Neutrophil migration depends on inflammatory chemokines and cytokines. To determine if blocking lymph flow impaired inflammatory chemokine/cytokine level in the LN, we sutured the lymphatic vessels before infection, and 2hpi. The contralateral LNs which were also infected and injected with Evans blue were collected as controls. All the LNs were collected at 4 hpi (**Figure 1C)**. Using multiplex chemokines/cytokines discovery array analysis, almost all of the chemokines related to neutrophil migration, such as CXCL1, CXCL2, and CCL2, as well as pro-inflammatory cytokines were suppressed when lymph flow was blocked either before infection or 2hpi (**Figure 1G, H and Table 1**).

Inflammatory chemokines and cytokines could be produced in the infected tissue and then travel with lymph to the LN. Lymph-borne bacteria may also stimulate LN resident cell to produce these factors. To distinguish these two sources of inflammatory chemokines and cytokines, we sutured lymphatic vessels of the pLNs before infection or 10 min after the infection. The contralateral LNs with infection and Evans blue injection were collected as control. All the LNs were collected at 4 hpi to quantify bacteria accumulation by counting CFUs (**Figure 1C)**. As expected, compared to the control pLNs, no *S. aureus* was detected in the LN when infected after suture. When lymphatic vessels were sutured 10 min post-infection, comparable numbers of *S. aureus* were detected in the pLNs (**Figure 1I**). In a separated study, using whole mount confocal imaging on the SCS, GFP labeled *S. aureus* was observed located in the SCS in both control mice and mice sutured 10 min post-infection, but not in the mice sutured before the infection (**Figure 1J**). Thus, bacteria had reached the LNs and were located in the SCS within 10 mins after the infection. Together, these results showed that without lymph flow, accumulation of *S. aureus* in the LN was not sufficient to activate inflammatory chemokine/cytokine production in the LN at 4hpi. At this time, lymph flow brought inflammatory chemokines from the infected tissue to the LN.

### Skin inflammation interrupted neutrophil positioning when exposed to a secondary infection

Previous reports showed that OX-skin inflammation suppresses lymphatic contraction and reduces lymph flow to the LN, which peaked at day 4 after OX treatment (OXd4) (37, 38, 42). To determine whether the interrupted lymph flow impacts neutrophil migration when exposed to a secondary infection with *S. aureus*, we collected the control or OXd4 LNs at 4hpi. To avoid the skin inflammation from impacting inflammatory responses at the site of infection, we sensitized skin at the abdomen and injected *S. aureus* at the intact skin in the flank (**Figure 2A**). OX-skin inflammation itself did not recruit a significant number of neutrophils, as seen in the contralateral LNs of the OXd4 LysM-GFP reporter mice (**Figure 2B**). In control LNs, neutrophils were found throughout the LN and were concentrated at the border of T and B cell zone, in the SCS and MS. In the OXd4 LNs, neutrophils appeared to be substantially reduced, and the remaining neutrophils were positioned in the MS (**Figure 2B**).

**Figure 2.**
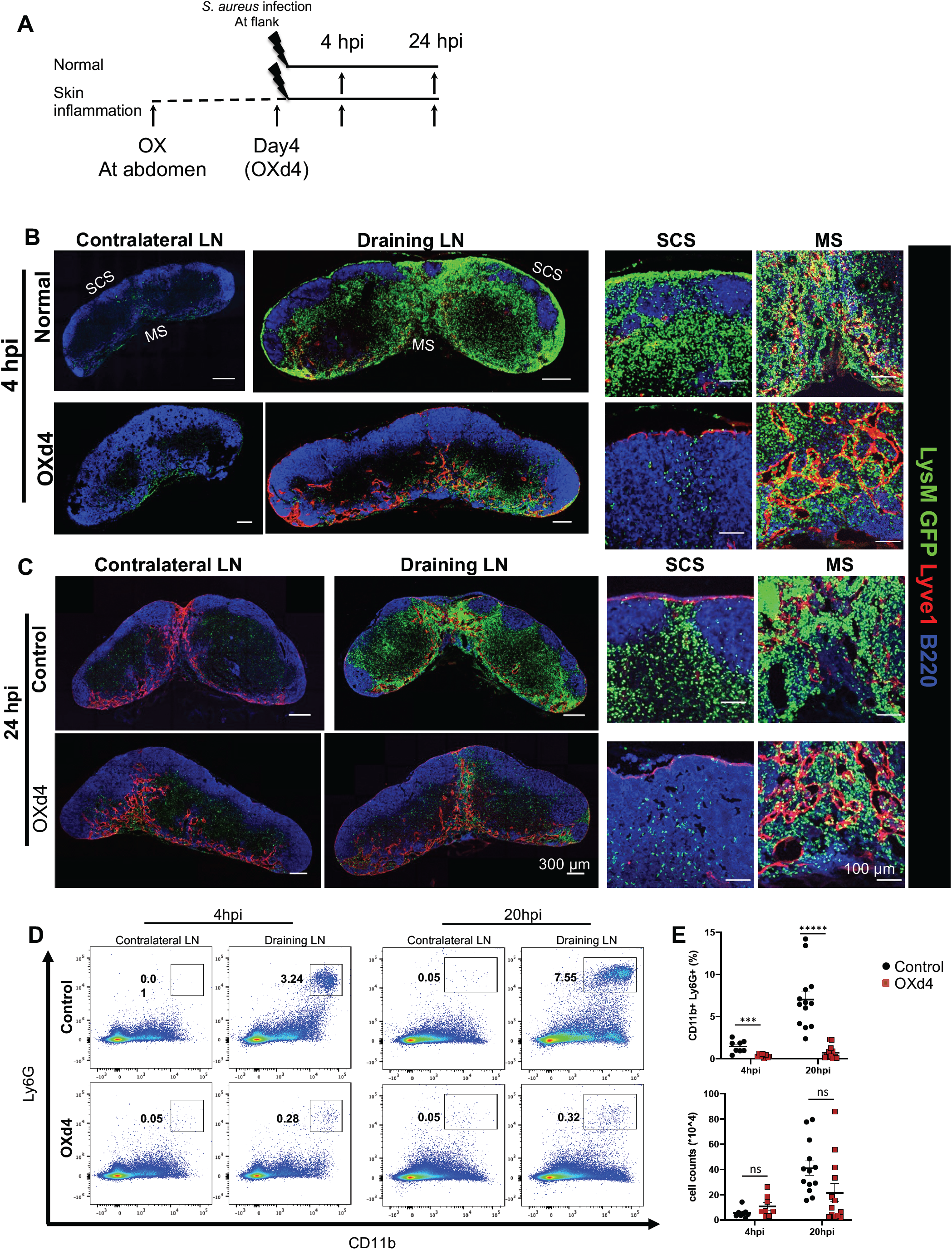
Inflammation interrupted neutrophil positioning when exposed to secondary infection. (**A**) The schematic diagram of the experimental design. Mice were treated with OX on their shaved abdomens. Four days after OX treatment (OXd4), control (uninflamed) and OXd4 (inflamed) mice were infected with 2.5×10^7^ *S. aureus* intradermally at the right flank. The contralateral and the draining inguinal LNs (iLNs) were collected at 4 hpi or 24hpi. (**B-C**) Using LysM-GFP reporter mice, cryosections were stained with anti-B220 (blue, B cell areas) and anti-Lyve1 (red, lymphatic vessel endothelial cells). Original magnification, 200×; 4hpi, n=4; 24hpi, n=3. (**D**) Using WT C57BL/6 mice, representative flow cytometry plot of neutrophils (CD11b^+^Ly6G^+^) in iLNs collected at 4hpi and 20 hpi. (**E**) Quantification of number of neutrophils per iLN collected at 4 and 20 hpi. Data shown as mean ±SEM, unpaired T-test; ***P < 0.001; ****P < 0.0005; ns, no significance; n=7-9.

To know if neutrophil recruitment or positioning in the SCS of the OXd4 LNs can be restored given a longer time, we collected the control and OXd4 LNs at 24hpi. By this time, in control LN, neutrophils had migrated away from the SCS. Most neutrophils were concentrated at the border of T and B cell zone, and in the MS. In the OXd4 LNs, neutrophils remained restricted to the MS, indicating that neutrophils could not migrate to the SCS even at a later time point (**Figure 2C**). Using WT mice, flow cytometry analysis showed that while the proportion of neutrophils (CD11b^+^Ly6G^+^) in the OXd4 LN was significantly reduced, the number of neutrophils per LN has no significant difference with the control LNs at either 4hpi or 20hpi because the total LN cell number had substantially expanded at OXd4 (**Figure 2D, E**). Therefore, when exposed to a secondary infection, neutrophil positioning, but not recruitment was impaired in the OX-inflamed LNs (OXd4).

### Neutrophil position in the SCS was independent of SCS macrophages

Next, we used multiplex chemokines/cytokines discovery array analysis to measure the levels of inflammatory chemokines in the LN when exposed to secondary infection at 4hpi. Without infection, chemokine/cytokine level was very low and was comparable between control and the OXd4 LNs. At 4hpi, inflammatory chemokines/cytokines were substantially increased in control LNs. However, the inflammatory chemokines were significantly lower in the OXd4 LNs (**Figure 3A, B and Table 2**). The question became why reduced chemokines in OXd4 LNs changed the positioning of neutrophil, but not their recruitment.

**Figure 3.**
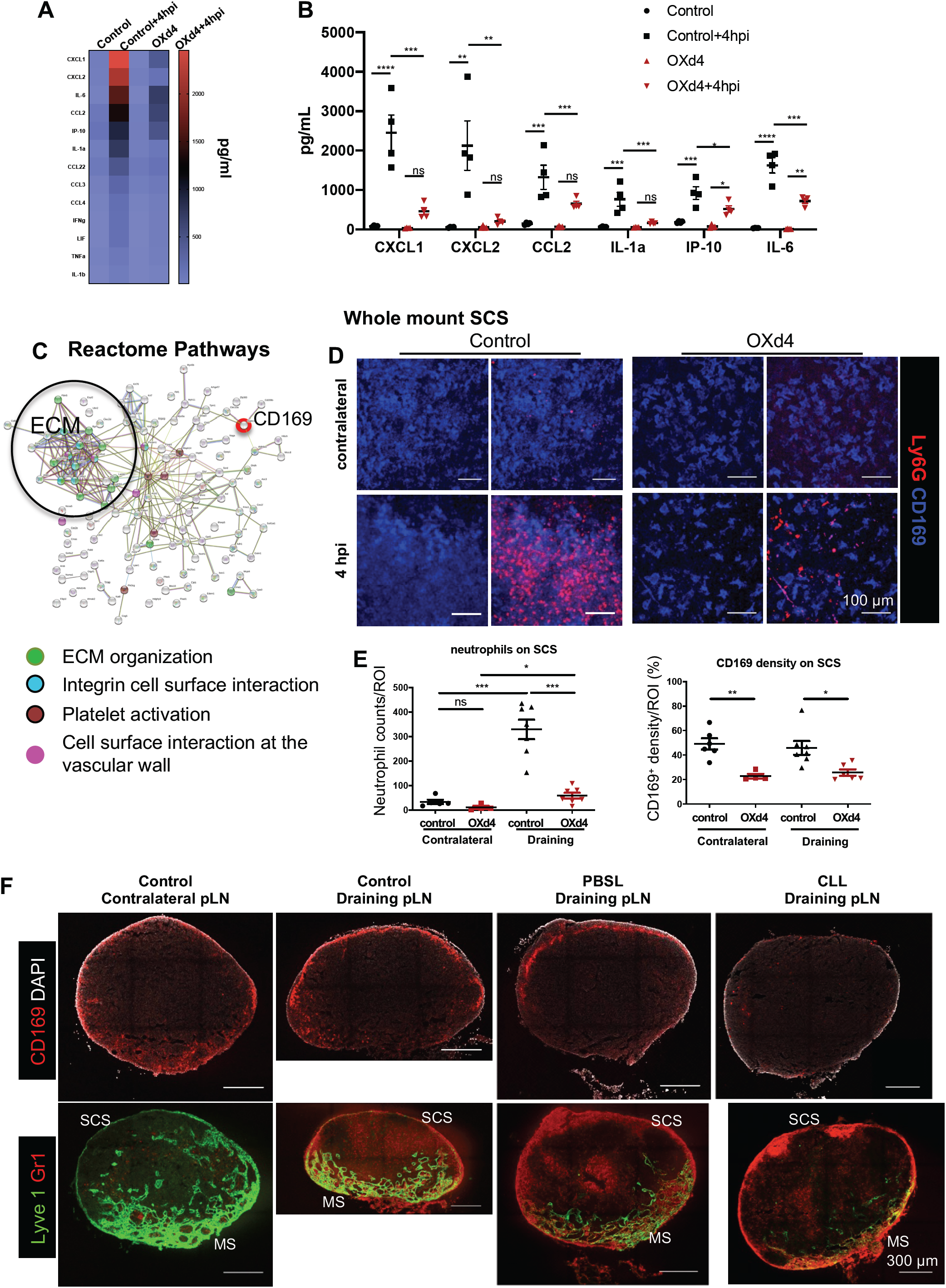
Neutrophil migration does not depend on SCS macrophages. (**A, B**) Control and OXd4 iLNs were harvested at 4hpi. The draining iLNs were homogenized in lysis buffer to extract proteins. (**A**) Heatmap analysis of cytokine/Chemokine discovery array. (**B**) Graph of Individual-values of high concentration cytokine/chemokine. Data shown as mean ±SEM, One-way ANOVA; *P<0.05; **P<0.01; ***P < 0.001; ****P < 0.0005; n=4. (**C**) Proteomics analysis. LNs were collected from control and OXd4 mice at 4hpi. Reactome pathway analysis using STRING-DB showed reduced ECM proteins and Siglec1(CD169) in the OXd4 LNs compared to the control LNs. (**D-E**) Using WT mice, iLN were collected from control and OXd4 mice at 4hpi. (**D**) Whole mount staining of iLN with anti-CD169 (blue, SCS macrophages) and anti-Ly6G (red) showed the distribution of neutrophils and SCS macrophages in the iLN SCS. Original magnification, 250×. (**E**) Quantification of neutrophil cell counts and SCS macrophage density per area of interest. n=7, Data shown as mean ±SEM, unpaired T-test; *P < 0.05; **P<0.01; ***P<0.001; ns, no significance. (**F**) PBS liposome (PBSL) or CLL was injected into the footpad of mice. At day 7 after liposome injection, 2.5×10^7^ *S. aureus* were injected into the footpad, and pLNs were collected 4hpi. Top panels, cryosections were stained with anti-CD169 (red) and DAPI (grey). Bottom panels, cryosections were stained with anti-Lyve1 (green) and anti-Gr-1 (red, neutrophils and monocytes). n=5 per group.

To better understand the molecular mechanism of how the inflammation changed neutrophil position in the OXd4 LNs, we collected control and OXd4 LNs at 4hpi to compare their proteomes (**Supplemental Figure 1**). Based on the Reactome Pathway analysis [using STRING v11.0 (https://string-db.org)] of the enriched proteins in control LNs at 4hpi, CD169 expression was significantly reduced in the OXd4 LNs (**Figure 3C**). Because CD169^+^ SCS macrophages are the first layer of cells in the LN encountering lymph-borne microbes and they play critical roles in bacterial phagocytosis and production of pro-inflammatory cytokines to recruit other types of cells, we hypothesized that the reduced CD169 macrophages impaired neutrophil migration at 4hpi in the OXd4 LN. Using WT mice, confocal images of whole-mount stained LNs showed that a large number of neutrophils (Ly6G^+^) and a dense layer of CD169^+^ SCS macrophages in the SCS at 4hpi in control LNs (**Figure 3D, left panel**). In OXd4 LNs, only a small number of neutrophils accumulated in the SCS, which was associated with a reduced SCS macrophage layer (**Figure 3C-E**). The reduction of the SCS macrophage layer in the OXd4 LNs was further confirmed using cryosections and flow cytometry analysis (**Supplemental Figure 1**). However, when used clodronate liposome (CLL) to deplete the LN macrophages, there was no difference in neutrophil recruitment or positioning in the SCS at 4hpi (**Figure 3F**). Therefore, neutrophil migration or positioning in the LN did not depend on SCS macrophages.

### LN remodeling impaired lymph flow along the conduits, but not around the sinus

Based on the Reactome Pathway analysis, we also identified a network of ECM proteins, including collagen I, II, IV and VI, as well as Laminin-1 and Laminin 2, that was reduced in the OXd4 LNs (**Figure 3A**). ECM proteins are major components of LN conduits, which facilitate lymph flow and traffic of small MW material from the SCS to the T cell zone and HEVs. Using immunofluorescent staining on cryosections, density and diameter of collagen I^+^ conduits were significantly reduced in the OXd4 LN compared to the control LNs (**Figure 4A and Supplemental Figure 2**). To understand how the reduced conduit density and diameter impacts lymph flow in the LN, control (non-OX treated) or OXd4 LNs were collected 2h after FITC treatment (**Figure 4B**). We detected much less FITC in LN conduits of the OXd4 LNs than those of control LNs (**Figure 4C**). To avoid the impacts of sample processing on FITC distribution, we used time-lapse intravital live imaging to track FITC distribution in LNs. Consistently, FITC entered the conduits and penetrated deep into the control LN (**Supplemental video 1**). In the OXd4 LNs, FITC was restricted to the sinus (**Supplemental video 2**).

**Figure 4.**
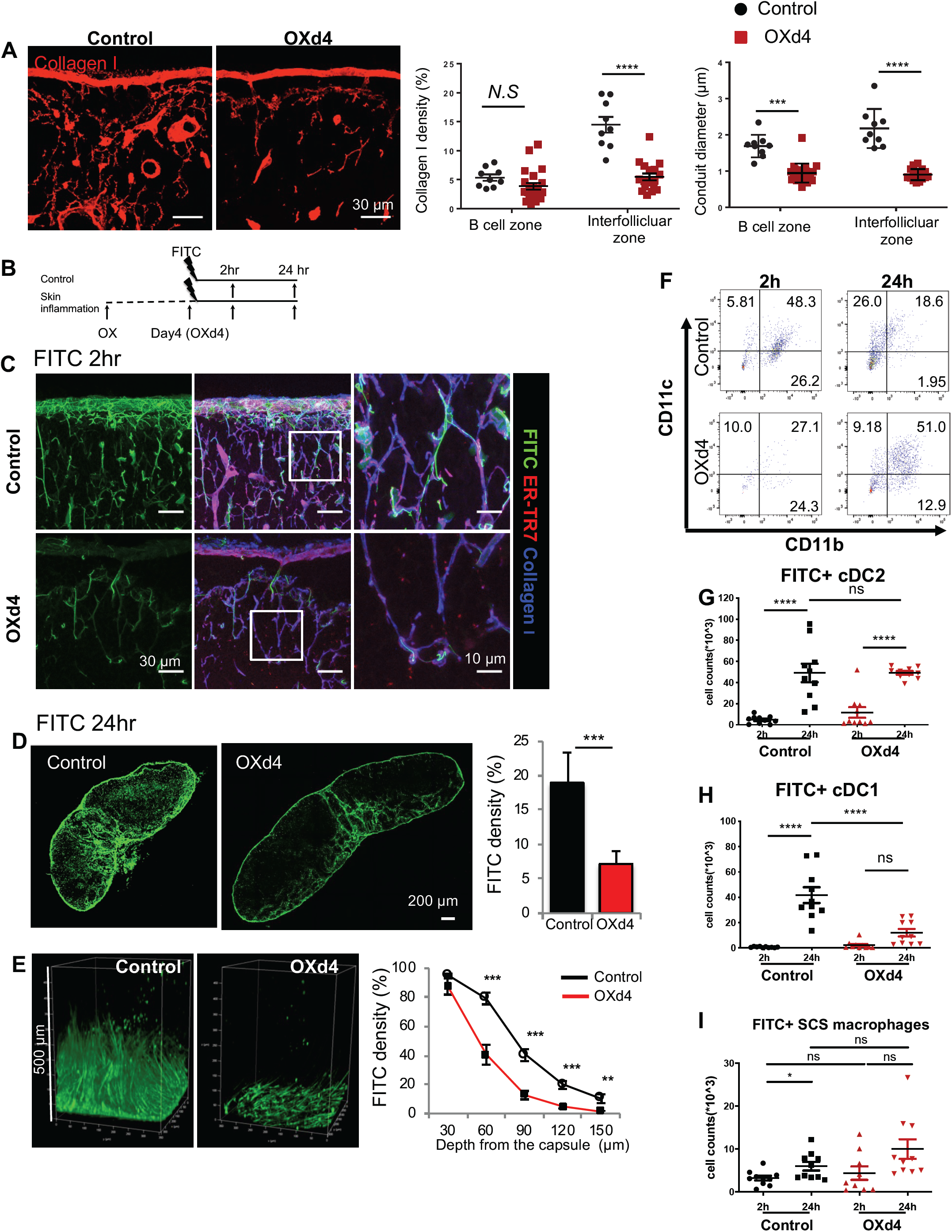
OX skin inflammation induced LN expansion impaired lymph flow along the conduits. (**A**) iLNs were harvested from control and OXd4 mice. Cryosections were stained with anti-collagen I antibody (red). The middle right and right panels are the quantifications of the density and diameter of the collagen I^+^ conduits in B cell zone and interfollicular zone, respectively, in control and OXd4 iLNs. Original magnification, 630×. Data shown as mean ±SEM, unpaired T-test; ***P < 0.001; ****P < 0.0001; ns, no significance; n=4-5 per group. (**B**) The schematic diagram of experimental design. Mice were treated with OX on the shaved abdomen. The control and OXd4 mice were treated with FITC at both flanks. After 2h or 24 h, the iLNs were collected for analysis. (**C**) FITC distribution with conduits in the iLNs at 2h after FITC sensitization. The cryosections were stained with anti-ERTR7 (red) and anti-collagen I (blue). Original magnification: left and middle columns, 630×; right column, 1890×. (**D**) The tile scan of cryosections (left and middle) and quantification of FITC density (right) were shown. The control and OXd4 iLNs were collected at 24h after FITC sensitization. Original magnification, 200×. Data shown as mean ±SEM, ***P < 0.001. (**E**) 3D-reconstructed images of whole mount LN showed FITC could not penetrate the LN efficiently in OXd4 LN. Control and OXd4 LNs were collected at 24h after FITC sensitization. The right chart showed FITC density correlated with depth from the capsule. Data shown as mean ±SEM, **P < 0.01; **P < 0.001; n>5 mice per group. **(F-I)** The capture of FITC by cDC1 but not cDC2 was interrupted in the OXd4 LNs. Control and OXd4 LNs were collected at 2h and 24h after FITC sensitization. (**F**) Representative flow cytometry plot of FITC population and sub-population of DCs. (**G-H**) FITC^+^cDC2 and cDC1 populations at 2h and 24h after FITC sensitization. (**I**) FITC^+^ SCS macrophage populations at 2h and 24h after FITC sensitization. Data shown as mean ±SEM, *P<0.05; **P < 0.01; ***P < 0.001; ns, no significance; n=10 iLNs/group.

Even at a more extended time point (24h after FITC treatment), FITC was still restricted to the SCS and MS in the OXd4 LNs compared to the control LNs (**Figure 4D**). By optical clearing with BABB, three dimensionally reconstructed images (3D) showed that FITC could not penetrate the OXd4 LNs as deep as the control LNs (**Figure 4E, left**). Quantification of FITC density every 30 μm from the LN SCS inward revealed that FITC density was comparable between control and the OXd4 LN at the first 30 μm, but it was substantially reduced in the OXd4 LNs for the subsequent 30-150 μm from the capsule (**Figure 4E, right**). Together, these data showed that OX-inflammation reduced conduits and impaired lymph flow along conduits but not in the SCS or MS.

It is known that from LN-resident dendritic DCs, cDC1 (CD11b^-^CD11c^+^) populations are located deeper in T cell zone and cDC2 (CD11b^+^CD11c^+^) populations are located close to the sinus (43). Flow cytometry results showed that the number of FITC^+^ cDC1 (DCs that captured FITC) were substantially reduced in OXd4 LNs compared to control LNs, while the number of FITC^+^ cDC2 and FITC^+^CD169^+^ SCS macrophages between control and OXd4 LNs at 2h and 24h after FITC sensitization are comparable (**Figure 4F-I**). These results further showed that lymph could reach the SCS and MS, but it did not effectively enter the T-cell zone in the OXd4 LNs.

### Inflammation impaired lymph reaching HEVs via conduits

Lymph-borne inflammatory chemokines should reach HEVs to direct neutrophil transmigration. Chemokines in lymph can diffuse from the sinuses to the nearby HEVs or flow along the conduits to the HEVs. Therefore, we compared lymph-HEV communication between control and OXd4 LNs. Using cryosections with IF staining, we measured the distance from the SCS or the MS to HEVs close to these regions in the LNs. The results showed that OX-skin inflammation substantially increased the distance from the SCS to HEVs, but not that of the MS to HEVs (**Figure 5A, B**). Using FITC as lymph tracer, at 4h after FITC treatment, a strong FITC signal was detected around HEVs, near both SCS and MS in control LNs (**Figure 5C, control**). In the OXd4 LNs, the FITC signal was mostly restricted to the sinuses. Only a small amount of FITC from the SCS could reach HEVs located in the interfollicular or T cell zone (close to SCS). In the MS, despite slightly weaker FITC signal, HEVs located in the MS remained exposed to FITC (**Figure 5C, OXd4**). These results suggested that due to the increased distance and reduced conduits in the OXd4 LNs, lymph preferentially reached HEVs located in the MS of the OXd4 LNs. Thus, lymph-borne chemokines that in OXd4 LNs should be able reach HEVs located in the MS, but could not effectively diffuse or flow along the conduits to reach HEVs in the interfollicular zone or in the T cell zone.

**Figure 5.**
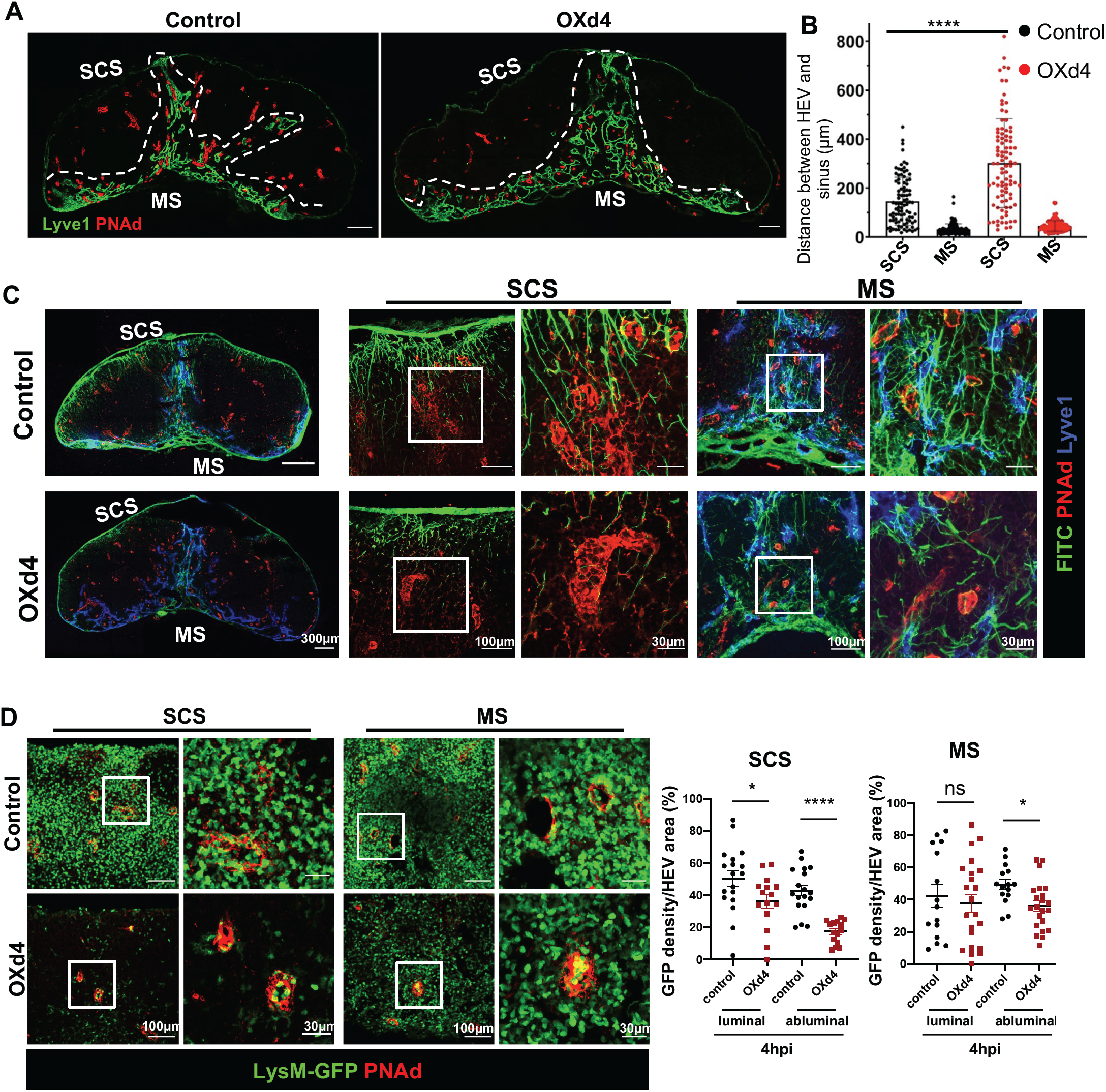
OX skin inflammation induced LN remodeling interrupted chemokine in lymph reach HEVs. (**A**) Distribution of HEVs in control and OXd4 LNs. iLNs were harvested from control and OXd4 mice and fixed with 4% PFA overnight. Cryosections were stained with anti-PNAd (red) and anti-Lyve1 (green) antibodies to show HEVs and sinuses. Original magnification, 200×; n=4. (**B**) The distance between HEVs with the SCS or MS in the normal and OXd4 LNs. Data shown as mean ±SEM; ****P < 0.0005; n=3-4. (**C**) FITC distribution in control and OXd4 iLNs. Control and OXd4 LNs were collected at 4h after FITC treatment. Cryosections were stained with anti-PNAd (red, HEVs) and anti-Lyve1(blue, lymphatic vessel endothelial cells) antibodies. (**D**) Neutrophil distribution around HEVs in control and OXd4 LN. Cryosections were stained with anti-PNAd (red, HEVs). Original magnification, 200×; zoomed-in images, 630×; n=3. Quantification of LysM-GFP cells intensity per HEV by ImageJ. GFP 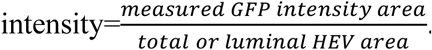Data shown as mean ±SEM, unpaired T-test; *P<0.05; **P<0.01; ****P < 0.0005; ns, no significance; n=4-5.

To determine if neutrophil preferentially migrate in HEVs in the MS due to the changed lymph flow direction, we used LysM-GFP mice to characterize neutrophil transmigration in HEVs located at different areas of LNs. In control LNs at 4hpi, neutrophils were detected in both the luminal and the abluminal side of HEVs across the LN (**Figure 5D, control**). In the OXd4 LNs, a slightly smaller number of neutrophils were detected in the lumen of the HEVs close to the SCS (interfollicular and the T cell zone). However, a substantially reduced neutrophil at the abluminal side of these HEVs indicated reduced neutrophil transmigration (**Figure 5D, SCS**). In the MS, there was no significant difference between the number of neutrophils in the lumen of HEVs compared to those in the control LNs. Neutrophils on the abluminal side of these HEVs were only slightly reduced compared to the control LNs (**Figure 5D, MS**). Collectively, neutrophils preferentially transmigrated at the HEVs located in the MS, where HEVs remained exposed to lymph (and lymph-borne chemokines) in the OXd4 LNs.

### Delayed bacterial clearance in the OXd4 LNs

Because bacteria were restricted in the SCS and neutrophil positioning in the SCS plays a critical role in phagocytosis and killing of the bacteria, we investigated if interrupted neutrophil positioning impaired *S. aureus* clearance in the OXd4 LN. Using LysM-GFP reporter mice, neutrophil positioning in the OXd4 pLNs was comparable with the iLNs (**Figure 6A**). We collected control or OXd4 pLNs at 4hpi and quantified the bacteria load by counting CFUs. The CFUs were not significantly different between control and OXd4 mice (**Figure 6B**). In a separate study using GFP-*S. aureus*, cryosections of LNs showed that GFP-*S. aureus* was restricted to the SCS in both control and OXd4 LNs (**Figure 6C**). Although bacteria appeared to be more scattered in the SCS of OXd4 pLNs due to LN cell expansion, the number and position of *S. aureus* were not changed in the OXd4 LNs at 4hpi.

**Figure 6.**
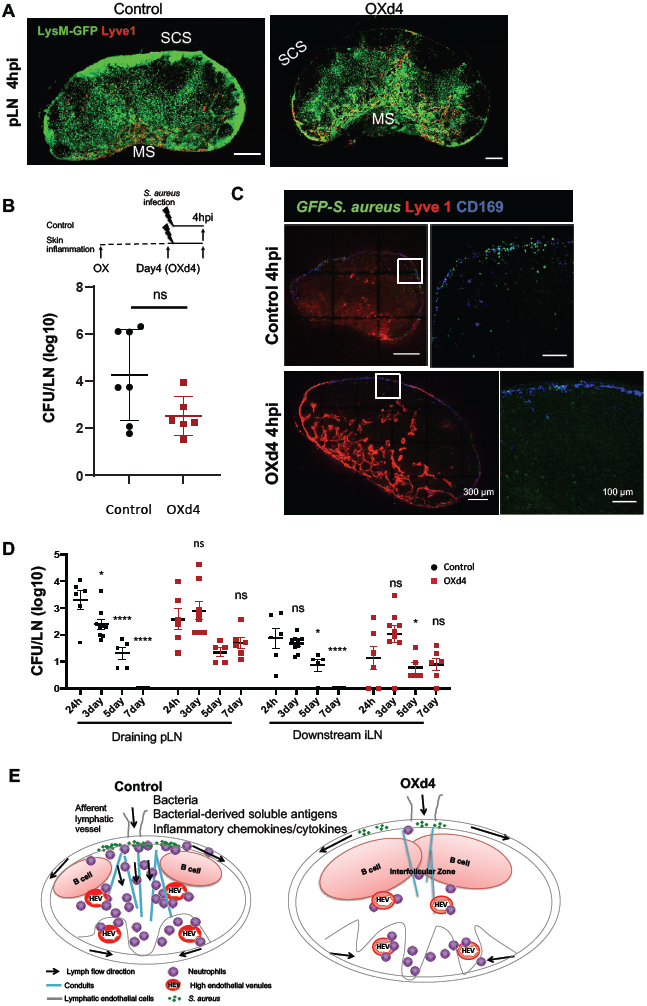
Delayed bacterial clearance in the OXd4 LNs. (**A**) Neutrophil distribution in pLNs. LysM-GFP mice were treated with OX on the leg, then 2.5⨯10^7^ *S. aureus* was injected intradermally at the footpad in control and OXd4 mice. The contralateral and draining pLNs were collected at 4hpi and fixed with 4% PFA overnight. Cryosections were stained with anti-Lyve-1 staining (red). (**B**) The schematic diagram of experimental design and CFU tests to quantify the number of *S. aureus. S. aureus* was injected in the footpad of control and OXd4 mice. The draining pLNs were collected at 4hpi. Data shown as mean ±SEM; ns, no significance. n=6 per group. (**C**) Bacteria distribution on the SCS in the control and OXd4 pLN. GFP*-S. aureus* was injected intradermally in the right footpad, and the draining pLNs were collected at 4hpi. Cryosections were stained with anti-Lyve1 (red) and anti-CD169 (blue, SCS macrophages). Original magnification, 200×; zoom in images, 630×. n=3. (**D**) Quantification of bacterial loads in control and OXd4 pLNs with *S. aureus* infection. The draining pLNs and downstream iLNs were collected at 24hpi, 3, 5, and 7 days post-infection. Data shown as mean ±SEM, *P<0.05; ns, no significance; n=6-8 per group. (**E**) Schematic diagram of neutrophil positioning in the LN. Soon after infection, lymph flow transports *S. aureus* to the SCS. Soluble antigens and soluble regulatory factors in lymph can reach the HEVs throughout the LN to direct neutrophils migration. Skin inflammation reduced inflammation chemokines in the LN. Furthermore, skin inflammation induced LN cell expansion and conduit remodeling. Lymph-borne chemokines could reach HEVs located in the MS but not those located in the interfollicular and T cell zone (close to the SCS). Consequently, neutrophils preferentially transmigrated through the HEVs in the MS and were positioned in the MS of the OXd4 LNs.

Finally, we quantified the numbers of bacterial at various time points, between 24hpi and 7 days post infection (dpi) to determine clearance of bacteria. In control pLNs, the bacterial load was substantially reduced from 24hpi to 3 dpi, and was entirely cleared by 7dpi. In OXd4 pLNs, the number of bacteria remained similar from 24hpi to 7 dpi. To determine if the changed lymph flow/neutrophil position caused spread of bacteria to the downstream LNs or into circulation, we also collected the downstream iLNs, liver and spleens to count the CFUs. Bacteria could spread to the downstream iLNs in both control and OXd4 mice by 24 hpi, but there was no significant difference between the number of *S. aureus* in the downstream iLNs, in control and OXd4 conditions at 24hpi (**Figure 6D**). No *S. aureus* was detected in the spleen or liver in control or OXd4 LNs (**data not shown**). Thus, there was no increase in the spread of bacteria to distant LNs or into the circulation during OX-skin inflammation. These results showed that impaired lymph flow was associated with altered neutrophil positioning and persistent infection in OXd4 LNs.

## Discussion

Skin draining LNs play a critical role in eliminating and preventing the systemic spread of pathogens during skin infection. One critical early step is the rapid recruitment of neutrophils through HEVs in the LNs (5, 6). By blocking lymph flow before or at a short time after infection, our studies showed that lymph flow is crucial for rapid recruitment and positioning of neutrophil in the LN after *S. aureus* infection. By 4hpi, lymphatic vessels transport bacteria and soluble factors (including bacterial-derived free-form antigens and infection-induced inflammatory chemokines and cytokines) from the skin to the LN. While *S. aureus* is restricted to the SCS, lymph flow (through the sinuses, along the conduits and by diffusion), facilitates lymph-borne soluble factor dispersal in the LN and direct neutrophil migration in HEVs first in the MS and then throughout the LN (**Figure 6E, control**).

Our study showed that blocking lymph flow soon after infection did not impact the number of *S. aureus* accumulation but globally reduced inflammatory chemokines and cytokines in the LN. Without lymph flow, the accumulation of *S. aureus* was insufficient to induce chemokine/cytokine production in the LN. These results suggested that most inflammatory cytokines/chemokines were produced in the infection site and then entered the LN with lymph flow. CD169^+^ SCS macrophages are the first layer of immune cells that encounter lymph-borne microbes in the LN. However, depletion of SCS macrophages suggested that neutrophil migration was independent of SCS macrophages. Together, activation of LN resident cells appeared to be non-essential for rapid neutrophil migration at 4hpi.

Skin inflammation induced lymphatic vessel and LN remodeling (OXd4). When exposed to a secondary infection, there was a global reduction of chemokine and cytokine in the OXd4 LNs. Due to the LN cell expansion and LN conduit reconstruction, lymph flow in the SCS could not effectively reach HEVs located in the interfollicular and T cell zone neither via diffusion nor conduits. Lymph-borne chemokines preferentially reached HEVs located in the MS, which resulted in most neutrophil migration occurring through the HEVs located in the MS (**Figure 6E**). Finally, reduced ECM proteins may impair the interaction with neutrophil surface integrins. From the proteomic analysis, integrin *α*1, *α*M, *β*1, *β*2, and *α*L were comparable between the control and OXd4 LNs (**Data not shown**). Thus, the changed migration is less likely caused by neutrophils surface integrins.

In summary, our study showed the importance of lymph flow in acute immune protection in infection. Interrupted lymph flow impaired neutrophil migration and resulted in persistent infection. The impaired lymph flow and neutrophil migration may contribute to the frequent infection in skin diseases, such as atopic dermatitis.

## Acknowledgment

The authors would like to thank Dr. Paul Kubes and Ania Bogoslowski for their critical input into this study. This work is supported by the University of Calgary start-up fund to SL, and the Kipnes Lymphatic Imaging Suite supported by the Dianne & Irving Kipnes Foundation. The Natural Sciences and Engineering Research Council of Canada (NSERC, 03641), Canadian Institute of Health Research (CIHR, PJT-156035) and Canada Foundation for Innovation (32930) to SL.

## Author contribution

Jingna Xue performed most of the experiments, data analysis and prepared manuscript. Yujia Lin, Darellynn Oo, Jianbo Zhang, Flavia Jesus, and Ava Zardynezhad performed some of the experiments. Luiz G. N. de Almeida, Daniel Young, and Antoine Dufour performed and analyzed the proteomic analysis. Shan Liao directed the project, performed some experiment, data analysis and prepared the manuscript.

## Conflict of Interest

The authors declare there is no conflict of interest.

---

**Supplementary Figure 1.**
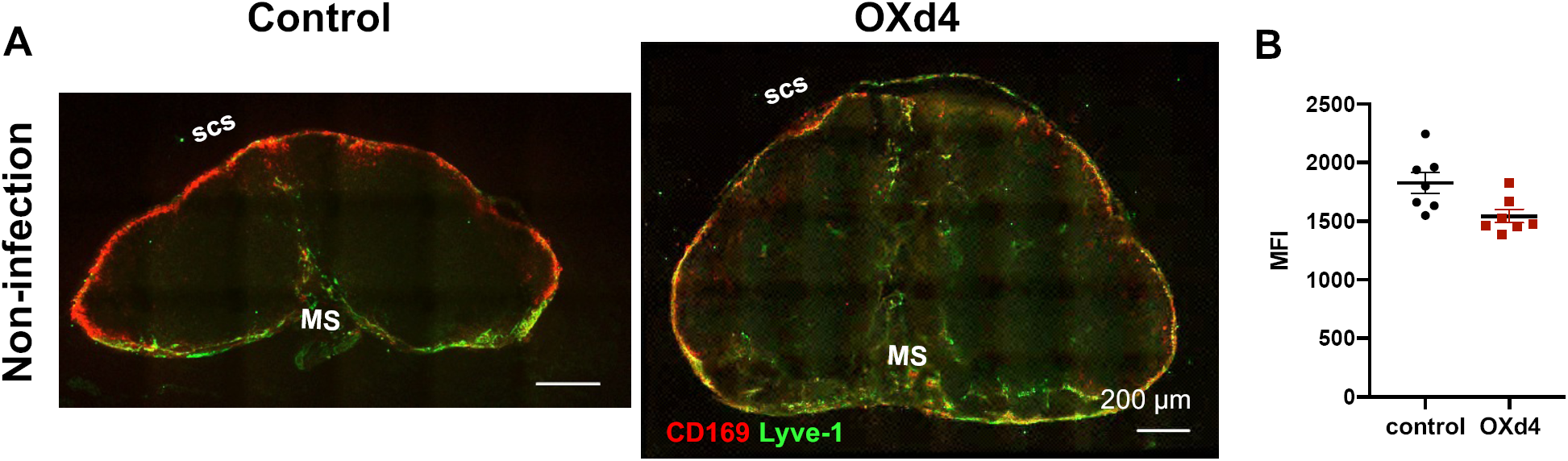
(**A**) Reduced CD169 macrophage layer in the LN SCS in OXd4 LN. iLNs were harvested from control and OXd4 mice. Cryosections were stained with anti-CD169 (red) and anti-Lyve1 (green). n=5. (**B**) FACS analysis showed reduced median fluorescent intensity (MFI) of CD169 in the OXd4 LNs. Data are mean ±SEM, unpaired T-test; *P<0.05; n=6.

**Supplementary Figure 2.**
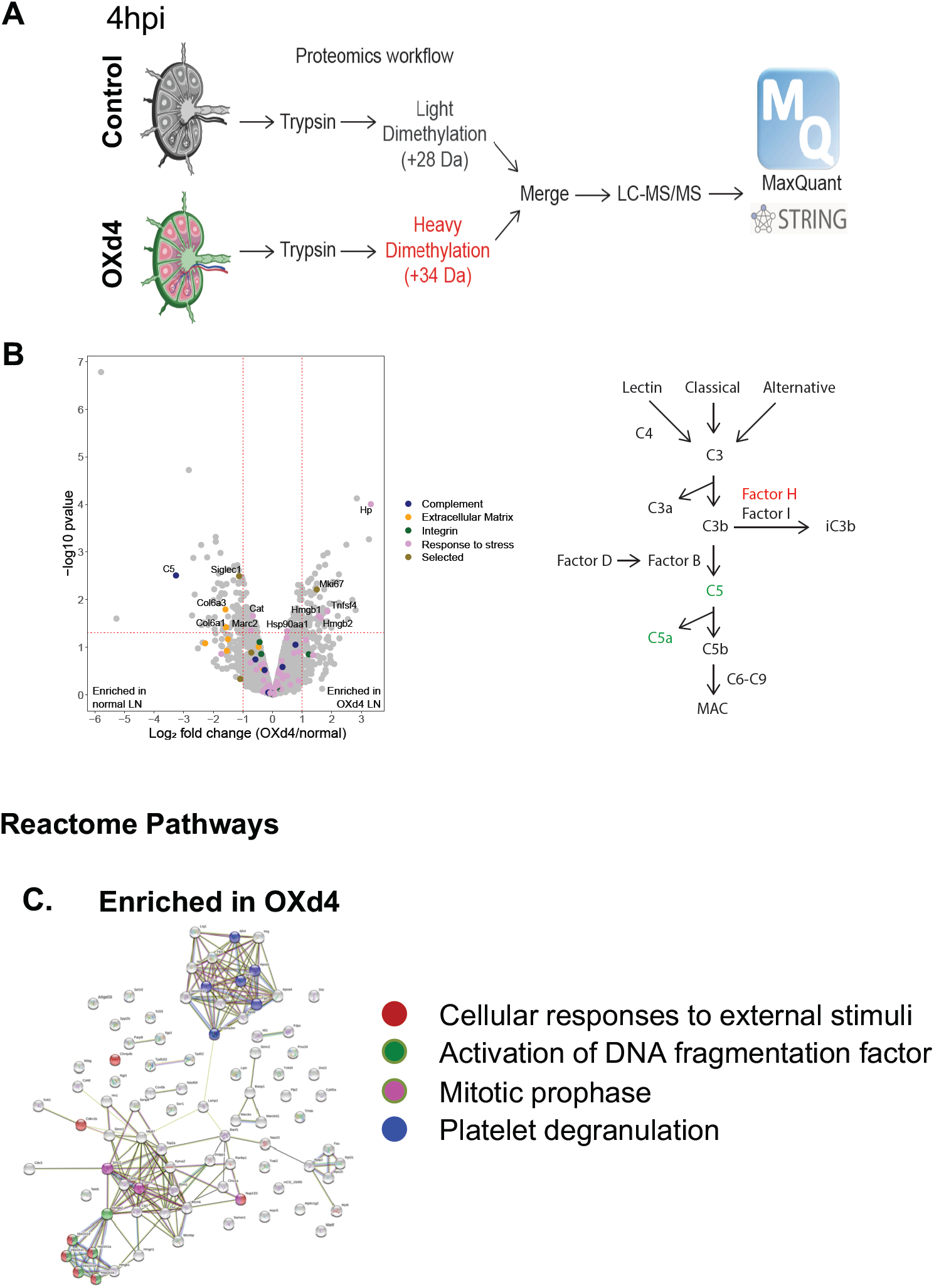
Proteomic analysis showed factors that were associated with proper neutrophil positioning. (A) Experimental Proteomics workflow of lymph nodes (n=4 per group). iLN were collected from control and OXd4 mice at 4hpi with *S. aureus* infection at the right flank. (B) *Left*, Volcano plot analysis of the proteomics data. *Right*, Diagram depicting a simplified complement cascade. Black, unchanged. Red, elevated in OXd4 LN. Green, elevated in control LN. Student t-test was used to determine statistical cutoffs. (C) Reactome pathway analysis using STRING-DB for proteins enriched in OXd4 LN.

**Supplementary Figure 3.**
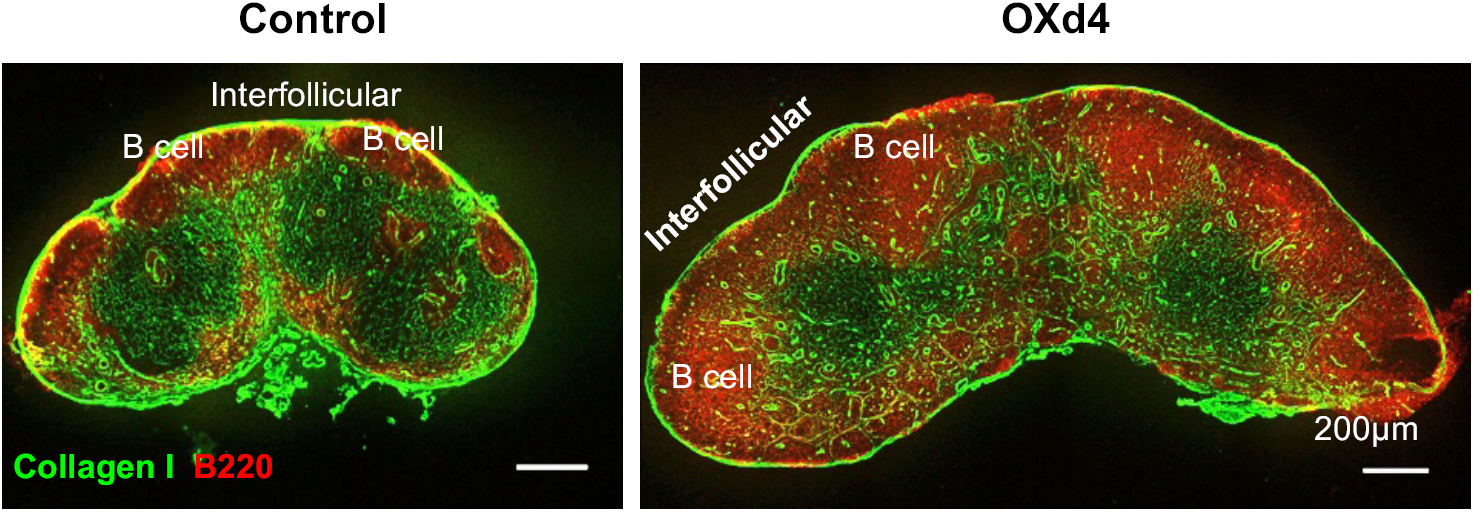
Conduit density was reduced in the OXd4 LN. iLNs were harvested from control and OXd4 mice. Cryosections were stained with anti-collagen I (green) and anti-B220 (red, B cells). B cell zone was dramatically expanded. Original magnification, 200×.

**Supplementary Table 1.** Multiplex chemokines/cytokines discovery array analysis when suture lymphatic vessels before and after infection.

|  | 4hpi |  |  | suture before |  |  | suture 2hpi |  |  |
| --- | --- | --- | --- | --- | --- | --- | --- | --- | --- |
|  | Mean | SEM | N | Mean | SEM | N | Mean | SEM | N |
| **CXCL2** | 3500.86 | 458.06756 | 4 | 119.8775 | 72.7688793 | 4 | 173.9075 | 25.8307738 | 4 |
| **IL-2** | 2443.2575 | 533.815517 | 4 | 670.785 | 214.252074 | 4 | 288.345 | 104.04897 | 4 |
| **MIG** | 1724.19 | 963.19314 | 4 | 322.1175 | 144.121663 | 4 | 159.8125 | 71.5267946 | 4 |
| **CXCL1** | 1334.7175 | 206.796048 | 4 | 79.2825 | 24.3955885 | 4 | 181.02 | 48.7679278 | 4 |
| **IL-6** | 1149.59 | 309.083651 | 4 | 52.88 | 22.4101227 | 4 | 76.2475 | 25.9821289 | 4 |
| **CCL2** | 1088.255 | 94.8172613 | 4 | 110.01 | 42.8214664 | 4 | 211.39 | 44.3380533 | 4 |
| **G-CSF** | 760.195 | 159.893208 | 4 | 69.1775 | 35.2468051 | 4 | 71.5625 | 25.2666098 | 4 |
| **IP-10** | 682.6225 | 93.2626262 | 4 | 220.9575 | 47.8461657 | 4 | 78.4275 | 22.0567988 | 4 |
| **IL-1a** | 407.2 | 54.0268879 | 4 | 37.575 | 8.96139638 | 4 | 48.5 | 11.8703707 | 4 |
| **Eotaxin** | 304.3575 | 78.5831282 | 4 | 62.68 | 14.1482042 | 4 | 128.28 | 43.943589 | 4 |
| **CCL4** | 272.34 | 36.9381088 | 4 | 11.055 | 8.45689216 | 4 | 25.5925 | 10.5120101 | 4 |
| **GM-CSF** | 197.6075 | 45.9603536 | 4 | 0.4675 | 0.4675 | 4 | 15.2933333 | 6.43088727 | 3 |
| **M-CSF** | 189.0975 | 63.8291846 | 4 | 13.31 | 1.89087722 | 4 | 80.235 | 17.3367598 | 4 |
| **CCL3** | 170.3275 | 42.5044347 | 4 | 1.74 | 1.74 | 4 | 16.1775 | 6.27004037 | 4 |
| **IL-17** | 137.7775 | 31.0595426 | 4 | 2.2775 | 1.20886569 | 4 | 12.3 | 1.28069643 | 4 |
| **LIF** | 117.8475 | 24.4493205 | 4 | 7.93 | 2.92095019 | 4 | 6.9025 | 1.54145156 | 4 |
| **IL-3** | 104.8425 | 27.3201764 | 4 | 5.2575 | 1.75979343 | 4 | 1.855 | 0.55354163 | 4 |
| **IFNg** | 73.28 | 18.7649416 | 4 | 2.3525 | 0.53457733 | 4 | 2.0825 | 0.60284568 | 4 |
| **IL-1b** | 52.0675 | 9.9459484 | 4 | 2.8625 | 0.65008173 | 4 | 4.1475 | 0.40594283 | 4 |
| **TNFa** | 48.5375 | 13.6514879 | 4 | 26.9125 | 6.95243527 | 4 | 27.4725 | 4.42073405 | 4 |
| **IL-4** | 11.5575 | 2.88722582 | 4 | 1.66 | 0.69112228 | 4 | 2.8925 | 0.92197591 | 4 |
| **IL-15** | 10.7675 | 2.04541917 | 4 | 8.0375 | 1.75896928 | 4 | 10.9575 | 0.34884034 | 4 |
| **RANTES** | 7.0875 | 0.95059083 | 4 | 3.1025 | 0.74202847 | 4 | 2.4925 | 0.34356889 | 4 |
| **VEGF** | 4.59 | 1.24661542 | 4 | 0.82 | 0.14124447 | 4 | 1.3575 | 0.10322911 | 4 |
| **IL-10** | 3.11 | 0.7732076 | 4 | 2.8325 | 0.72720212 | 4 | 1.67 | 0.53915057 | 4 |
| **IL-9** | 2.9825 | 0.97522967 | 4 | 4.2125 | 0.90982027 | 4 | 4.83 | 1.50919957 | 4 |
| **IL-7** | 2.925 | 0.88228397 | 4 | 5.7575 | 0.54696702 | 4 | 8.5375 | 1.62585554 | 4 |
| **IL-5** | ND |  |  | ND |  |  | ND |  |  |
| **IL-12p40** | ND |  |  | ND |  |  | ND |  |  |
| **IL-12p70** | ND |  |  | ND |  |  | ND |  |  |
| **LIX** | ND |  |  | ND |  |  | ND |  |  |

**Supplementary Table 2.** Multiplex chemokines/cytokines discovery array analysis in control and OXd4 LNs

|  | Control |  |  | Control+4hpi |  |  | OXd4 |  |  | OXd4+4hpi |  |  |
| --- | --- | --- | --- | --- | --- | --- | --- | --- | --- | --- | --- | --- |
|  | Mean | SEM | N | Mean | SEM | N | Mean | SEM | N | Mean | SEM | N |
| **6Ckine/Exodus 2** | 9696.02 | 2944.57 | 2 | 20382.84 | 2189.84 | 3 | 1608183.82 | 1597958.17 | 3 | 71160.50 | 33523.45 | 4 |
| **MCP-5** | 4844.10 | 1772.39 | 4 | 4947.25 | 1676.65 | 4 | 1181.54 | 126.11 | 4 | 7020.55 | 981.04 | 4 |
| **IL-2** | 205.34 | 11.58 | 4 | 2498.15 | 1055.18 | 3 | 52.43 | 5.63 | 4 | 436.98 | 46.50 | 4 |
| **CXCL1** | 84.13 | 8.67 | 4 | 2456.58 | 448.67 | 4 | 38.96 | 4.62 | 4 | 463.09 | 95.65 | 4 |
| **TIMP-1** | 193.88 | 27.18 | 4 | 2139.84 | 525.39 | 4 | 1381.74 | 517.75 | 4 | 2155.48 | 285.12 | 4 |
| **CXCL2** | 58.82 | 7.16 | 4 | 2128.72 | 628.14 | 4 | 61.58 | 15.65 | 4 | 208.11 | 44.16 | 4 |
| **IL-6** | 38.68 | 3.34 | 4 | 1627.54 | 190.06 | 4 | 15.39 | 2.50 | 4 | 724.73 | 61.00 | 4 |
| **CCL2** | 147.28 | 14.19 | 4 | 1325.40 | 306.15 | 4 | 71.66 | 10.04 | 4 | 651.72 | 67.52 | 4 |
| **IP-10** | 188.80 | 12.67 | 4 | 925.04 | 159.54 | 4 | 78.34 | 21.86 | 4 | 522.32 | 83.84 | 4 |
| **IL-1a** | 68.05 | 5.28 | 4 | 767.22 | 178.06 | 4 | 58.14 | 3.41 | 4 | 163.92 | 12.54 | 4 |
| **CCL22** | 130.41 | 26.81 | 3 | 499.05 | 113.35 | 3 | 126.46 | 5.30 | 3 | 200.59 | 9.57 | 3 |
| **IL-11** | 0.19 | 0.19 | 4 | 367.60 | 124.17 | 4 | 0.48 | 0.48 | 4 | 193.69 | 60.36 | 4 |
| **Eotaxin** | 111.40 | 10.94 | 4 | 364.85 | 41.92 | 4 | 76.28 | 7.45 | 4 | 207.29 | 35.91 | 4 |
| **G-CSF** | 41.72 | 4.88 | 4 | 354.94 | 149.60 | 4 | 25.88 | 3.00 | 4 | 241.55 | 123.16 | 4 |
| **CCL3** | 52.02 | 3.84 | 4 | 323.29 | 57.54 | 4 | 70.02 | 4.35 | 4 | 120.86 | 8.99 | 4 |
| **IL-3** | 4.62 | 0.46 | 4 | 285.22 | 65.49 | 4 | 4.85 | 1.02 | 4 | 193.19 | 49.61 | 4 |
| **IL-17** | 1.26 | 0.28 | 4 | 273.77 | 36.66 | 4 | 0.64 | 0.15 | 3 | 19.76 | 5.11 | 4 |
| **CCL4** | 28.15 | 3.05 | 4 | 239.50 | 48.41 | 4 | 23.38 | 2.16 | 3 | 83.28 | 5.01 | 4 |
| **IFNg** | 11.74 | 1.27 | 4 | 230.45 | 66.20 | 4 | 7.71 | 0.86 | 4 | 77.11 | 8.37 | 4 |
| **LIF** | 8.57 | 2.70 | 4 | 221.83 | 43.63 | 4 | 6.81 | 0.86 | 4 | 83.95 | 19.13 | 4 |
| **GM-CSF** | 36.37 | 1.93 | 4 | 193.96 | 31.32 | 4 | 28.14 | 2.80 | 3 | 107.58 | 12.51 | 4 |
| **TNFa** | 33.10 | 6.08 | 3 | 135.31 | 22.22 | 4 | 17.76 | 1.73 | 4 | 61.58 | 6.19 | 4 |
| **CCL19** | 75.46 | 11.86 | 4 | 117.99 | 10.61 | 4 | 115.66 | 31.12 | 4 | 119.47 | 8.48 | 4 |
| **IL-1b** | 7.96 | 0.58 | 3 | 94.94 | 30.50 | 4 | 11.02 | 0.44 | 4 | 39.93 | 4.90 | 4 |
| **LIX** | 17.66 | 1.56 | 4 | 73.45 | 10.31 | 4 | 27.41 | 3.57 | 4 | 60.58 | 8.57 | 4 |
| **IL-15** | 26.12 | 7.30 | 4 | 46.04 | 9.74 | 4 | 25.49 | 3.10 | 4 | 29.77 | 6.30 | 4 |
| **M-CSF** | 16.80 | 3.07 | 4 | 40.32 | 7.42 | 4 | 13.85 | 1.33 | 4 | 23.59 | 2.80 | 4 |
| **CX3CL1** | 16.94 | 3.37 | 4 | 29.35 | 6.26 | 4 | 8.75 | 2.55 | 4 | 13.61 | 2.09 | 4 |
| **IL-12(p70)** | 11.02 | 1.58 | 4 | 19.81 | 5.08 | 4 | 10.55 | 1.89 | 4 | 13.22 | 2.13 | 4 |
| **IL-7** | 20.15 | 4.48 | 4 | 15.84 | 1.55 | 4 | 20.67 | 2.19 | 4 | 17.38 | 3.71 | 4 |
| **IL-13** | 6.49 | 1.53 | 4 | 14.01 | 2.14 | 4 | 8.81 | 1.36 | 4 | 14.46 | 2.46 | 4 |
| **RANTES** | 10.12 | 2.53 | 4 | 13.85 | 1.74 | 4 | 6.33 | 0.34 | 4 | 9.60 | 0.49 | 4 |
| **IL-9** | 30.86 | 5.81 | 4 | 13.69 | 0.81 | 4 | 20.33 | 2.09 | 4 | 20.13 | 1.77 | 4 |
| **IL-12(p40)** | 7.37 | 1.50 | 4 | 11.94 | 3.34 | 4 | 7.38 | 0.93 | 4 | 10.49 | 1.52 | 4 |
| **VEGF** | 2.29 | 0.33 | 4 | 11.34 | 0.83 | 4 | 2.25 | 0.12 | 4 | 3.68 | 0.13 | 4 |
| **IL-5** | 3.10 | 0.23 | 4 | 10.71 | 2.06 | 4 | 2.28 | 0.33 | 4 | 7.80 | 1.65 | 4 |
| **IL-10** | 3.14 | 0.64 | 4 | 5.74 | 0.88 | 4 | 4.33 | 0.63 | 4 | 6.25 | 0.91 | 4 |
| **CCL20** | 3.23 | 0.71 | 4 | 4.35 | 1.03 | 4 | 4.55 | 0.44 | 4 | 11.87 | 1.08 | 4 |
| **EPO** | ND |  |  | ND |  |  | ND |  |  | ND |  |  |
| **IFNb-1** | ND |  |  | ND |  |  | ND |  |  | ND |  |  |
| **IL-20** | ND |  |  | ND |  |  | ND |  |  | ND |  |  |

**Supplemental Video 1**. *Intravital* live imaging of FITC trafficking in the control LN. After LNs were surgically exposed and set on stage, 5 μl of FITC was injected at the footpad. Approximately 30 min later, FITC had filled most of the conduits in the LN. The video started at around 30 minutes at FITC injection, and time-lapse was taken at 20 seconds/frame for 20-30 minutes.

**Supplemental Video 2**. *Intravital* live imaging of FITC trafficking in the OXd4 LN. Same as in the control mice, the video started at approximately 30 minutes at FITC injection, time-lapse were taken at 20 seconds/frame for 20-30 minutes.

## Notes

### Competing Interest Statement

The authors have declared no competing interest.

## Reference

1. W. Kastenmuller, P. Torabi-Parizi, N. Subramanian, T. Lammermann, R. N. Germain, A spatially-organized multicellular innate immune response in lymph nodes limits systemic pathogen spread. Cell 150, 1235–1248 (2012).

2. M. Y. Gerner, P. Torabi-Parizi, R. N. Germain, Strategically localized dendritic cells promote rapid T cell responses to lymph-borne particulate antigens. Immunity 42, 172–185 (2015).

3. S. Liao, P. Y. von der Weid, Lymphatic system: An active pathway for immune protection. Semin Cell Dev Biol (2014) https:/doi.org/10.1016/j.semcdb.2014.11.012.

4. H. Qi, W. Kastenmüller, R. N. Germain, Spatiotemporal Basis of Innate and Adaptive Immunity in Secondary Lymphoid Tissue. Annu. Rev. Cell Dev. Biol. 30, 141–167 (2014).

5. T. Chtanova, et al., Dynamics of neutrophil migration in lymph nodes during infection. Immunity 29, 487–496 (2008).

6. A. Bogoslowski, E. C. Butcher, P. Kubes, Neutrophils recruited through high endothelial venules of the lymph nodes via PNAd intercept disseminating *Staphylococcus aureus*. Proc. Natl. Acad. Sci. 115, 201715756 (2018).

7. O. Kamenyeva, et al., Neutrophil Recruitment to Lymph Nodes Limits Local Humoral Response to Staphylococcus aureus. PLoS Pathog. 11 (2015).

8. K. Brulois, et al., A molecular map of murine lymph node blood vascular endothelium at single cell resolution. Nat. Commun. (2020) https:/doi.org/10.1038/s41467-020-17291-5.

9. U. H. von Andrian, C. M’Rini, In situ analysis of lymphocyte migration to lymph nodes. Cell Adhes Commun 6, 85–96 (1998).

10. A. O. Anderson, N. D. Anderson, Studies on the structure and permeability of the microvasculature in normal rat lymph nodes. Am J Pathol 80, 387–418 (1975).

11. D. L. Drayton, S. Liao, R. H. Mounzer, N. H. Ruddle, Lymphoid organ development: From ontogeny to neogenesis. Nat. Immunol. 7, 344–353 (2006).

12. A. B. Raff, D. Kroshinsky, Cellulitis: A Review. JAMA 316, 325–337 (2016).

13. T. Junt, et al., Subcapsular sinus macrophages in lymph nodes clear lymph-borne viruses and present them to antiviral B cells. Nature 450, 110–114 (2007).

14. K. Asano, et al., CD169-Positive Macrophages Dominate Antitumor Immunity by Crosspresenting Dead Cell-Associated Antigens. Immunity 34, 85–95 (2011).

15. L. Martinez-Pomares, S. Gordon, CD169^+^ macrophages at the crossroads of antigen presentation. Trends Immunol. 33, 66–70 (2012).

16. B. Ravishankar, et al., Marginal zone CD169+ macrophages coordinate apoptotic cell-driven cellular recruitment and tolerance. Proc. Natl. Acad. Sci. U. S. A. 111, 4215–4220 (2014).

17. F. Pucci, et al., SCS macrophages suppress melanoma by restricting tumor-derived vesicle{\textendash}B cell interactions. Science (80-.). 352, 242–246 (2016).

18. M. Y. Gerner, P. Torabi-Parizi, R. N. Germain, Strategically Localized Dendritic Cells Promote Rapid T Cell Responses to Lymph-Borne Particulate Antigens. Immunity 42, 172–185 (2015).

19. M. Iannacone, et al., Subcapsular sinus macrophages prevent CNS invasion on peripheral infection with a neurotropic virus. Nature 465, 1079–1083 (2010).

20. D. A. P. Louie, S. Liao, Lymph Node Subcapsular Sinus Macrophages as the Frontline of Lymphatic Immune Defense. Front. Immunol. 10, 347 (2019).

21. M. Y. Gerner, K. A. Casey, W. Kastenmuller, R. N. Germain, Dendritic cell and antigen dispersal landscapes regulate T cell immunity. J. Exp. Med. 214, 3105–3122 (2017).

22. J. E. Chang, S. J. Turley, Stromal infrastructure of the lymph node and coordination of immunity. Trends Immunol 36, 30–39 (2015).

23. R. Roozendaal, et al., Conduits Mediate Transport of Low Molecular Weight Antigen to Lymph Node Follicles. Immunity 30, 264–276 (2009).

24. M. Sixt, et al., The conduit system transports soluble antigens from the afferent lymph to resident dendritic cells in the T cell area of the lymph node. Immunity 22, 19–29 (2005).

25. M. Jafarnejad, M. C. Woodruff, D. C. Zawieja, M. C. Carroll, J. E. Moore, Modeling lymph flow and fluid exchange with blood vessels in lymph nodes. Lymphat. Res. Biol. 13, 234–247 (2015).

26. J. E. Gretz, C. C. Norbury, A. O. Anderson, A. E. Proudfoot, S. Shaw, Lymph-borne chemokines and other low molecular weight molecules reach high endothelial venules via specialized conduits while a functional barrier limits access to the lymphocyte microenvironments in lymph node cortex. J Exp Med 192, 1425–1440 (2000).

27. R. E. Mebius, P. R. Streeter, J. Breve, A. M. Duijvestijn, G. Kraal, The influence of afferent lymphatic vessel interruption on vascular addressin expression. J Cell Biol 115, 85–95 (1991).

28. R. E. Mebius, et al., Expression of GlyCAM-1, an endothelial ligand for L-selectin, is affected by afferent lymphatic flow. J Immunol 151, 6769–6776 (1993).

29. A. A. Tomei, S. Siegert, M. R. Britschgi, S. A. Luther, M. A. Swartz, Fluid Flow Regulates Stromal Cell Organization and CCL21 Expression in a Tissue-Engineered Lymph Node Microenvironment. J. Immunol. 183, 4273–4283 (2009).

30. J. E. Chang, M. B. Buechler, E. Gressier, S. J. Turley, M. C. Carroll, Mechanosensing by Peyer’s patch stroma regulates lymphocyte migration and mucosal antibody responses. Nat. Immunol. 20, 1506–1516 (2019).

31. S. E. Acton, et al., Dendritic cells control fibroblastic reticular network tension and lymph node expansion. Nature 514, 498–502 (2014).

32. V. G. Martinez, et al., Fibroblastic Reticular Cells Control Conduit Matrix Deposition during Lymph Node Expansion. Cell Rep 29, 2810–2822 e5 (2019).

33. R. E. Mebius, J. Breve, A. M. Duijvestijn, G. Kraal, The function of high endothelial venules in mouse lymph nodes stimulated by oxazolone. Immunology 71, 423–427 (1990).

34. D. Hoke, et al., Selective modulation of the expression of L-selectin ligands by an immune response. Curr. Biol. 5, 670–678 (1995).

35. V. V Swarte, et al., Regulation of fucosyltransferase-VII expression in peripheral lymph node high endothelial venules. Eur J Immunol 28, 3040–3047 (1998).

36. K. Veerman, C. Tardiveau, F. Martins, J. Coudert, J. P. Girard, Single-Cell Analysis Reveals Heterogeneity of High Endothelial Venules and Different Regulation of Genes Controlling Lymphocyte Entry to Lymph Nodes. Cell Rep 26, 3116–3131 e5 (2019).

37. S. Liao, N. H. Ruddle, Synchrony of High Endothelial Venules and Lymphatic Vessels Revealed by Immunization. J. Immunol. 177, 3369–3379 (2006).

38. S. Liao, et al., Impaired lymphatic contraction associated with immunosuppression. Proc Natl Acad Sci U S A 108, 18784–18789 (2011).

39. Y. Lin, J. Xue, S. Liao, Blocking Lymph Flow by Suturing Afferent Lymphatic Vessels in Mice. J. Visulized Exp., 1–5 (2020).

40. Y. Lin, et al., Perinodal Adipose Tissue Participates in Immune Protection through a Lymphatic Vessel–Independent Route. J. Immunol. 201, 296–305 (2018).

41. M. T.R., H. S.E., V. A. U.H., T-cell priming by dendritic cells in lymph nodes occurs in three distinct phases. Nature 427, 154–159 (2004).

42. D. Aebischer, A. H. Willrodt, C. Halin, Oxazolone-induced contact hypersensitivity reduces lymphatic drainage but enhances the induction of adaptive immunity. PLoS One 9, e99297 (2014).

43. M. Y. Gerner, K. A. Casey, W. Kastenmuller, R. N. Germain, Dendritic cell and antigen dispersal landscapes regulate T cell immunity. J Exp Med (2017) https:/doi.org/10.1084/jem.20170335.

